# Giving it a fair shake: A simple, fast, & efficient method to extract amatoxins from the death cap mushroom, *Amanita phalloides*

**DOI:** 10.1101/2024.11.15.623858

**Authors:** Catharine A. Adams, Candace S. Bever, Jacquelyn M. Blake-Hedges, Sean Brown, Mitchell G. Thompson, Scott Behie, Jay Keasling, Patrick Shih

**Affiliations:** Department of Plant and Microbial Biology, University of California-Berkeley, Berkeley, California, USA; Environmental Genomics and Systems Biology Division, Lawrence Berkeley National Laboratory, Berkeley, California, USA; Foodborne Toxin Detection and Prevention Research Unit, Western Regional Research Center, United States Department of Agriculture, Agricultural Research Service, Albany, CA, USA; Department of Chemistry, University of California-Berkeley, Berkeley, CA, USA; Joint BioEnergy Institute, Emeryville, CA, USA; Department of Bioengineering, University of California, Berkeley, California, USA; Biological Systems & Engineering Division, Lawrence Berkeley National Laboratory, Berkeley, CA 94720, USA; Department of Chemical and Biomolecular Engineering, University of California, Berkeley, CA 94720, USA; QB3, University of California, Berkeley, Berkeley, CA, USA; Center for Biosustainability, Danish Technical University; Innovative Genomics Institute, Berkeley, California, USA

## Abstract

The death cap mushroom, *Amanita phalloides*, is well known for containing amatoxins such as alpha- and beta-amanitin, which inhibit eukaryotic RNA polymerase II. While these toxins have been used in research for almost a century, they have recently garnered attention for their role in drug-antibody conjugates. The amatoxins are still largely extracted from wild mushrooms, which cannot be made to fruit in the lab.

We propose simplified extraction methods that could reduce hazardous exposures to dust and expedite sample analysis without sacrificing accuracy. We recently developed a Lateral Flow Immunoassay (LFIA), for which we identified that sample maceration was not needed to extract the amatoxins and that the incubation time for extraction could be accomplished in 1 minute. In this current work, we hypothesized that these same extraction adjustments–minimal tissue maceration and reduced incubation time–could be transferable to instrumental detection methods.

To test the need for sample maceration, we utilized three different techniques: 1) traditional mortar and pestle, 2) a similarly disruptive method of bead beating, and 3) no grinding, but rather hand shaking dried mushroom tissue in extraction buffer. In addition, we performed the solvent extraction step at varying times to observe if more time allows for more toxin to be removed from the tissue. Lastly, we utilized two comparable solvent evaporation methods (rotovap or speedvac) to establish if multiple samples could be processed simultaneously, thus improving sample throughput.

We adjusted aspects of the typical extraction protocol, which resulted in a rapid (1 min) incubation step, along with minimal sample handling (no grinding) of the dried mushroom tissue. We present an extraction protocol that saves time, reduces equipment contamination, and minimizes risk to the researcher. The impact of this faster, safer method may help produce these important toxins faster, for both research and medical use.

## 1.2. Introduction

Wild mushrooms produce many compounds of biological significance (Lull et al. 2005; Holliday and Cleaver 2008; Jayakumar et al. 2008; Heleno et al. 2011; Alves et al. 2012; Heleno et al. 2013; Soares et al. 2013; Chang et al. 2015; Kozarski et al. 2015; Phan et al. 2015; Taofiq et al. 2016; Daley et al. 2017; Wasser 2017). One mushroom species of intense pharmacological interest is *Amanita phalloides*, also known as the death cap mushroom. The death cap is well known for producing the bicyclic octapeptides called amatoxins which inhibit RNA polymerase II (Wieland 1983; Carter and Drouin 2009), as well as the related heptapeptides the phallotoxins which inhibit actin polymerization (Wieland and Govindan 1974; Vandekerckhove et al. 1985). Both toxins have been used in cell research for decades (e.g. (Schultz and Hall 1976; Wieland and Faulstich 1978; Warn and Magrath 1983; Jendrisak 1980; Anderl et al. 2012). Furthermore, the primary amatoxin, alpha-amanitin, has been found to have strong anti-tumor effects. It was effective against the MCF-7 breast cancer cell line (Kaya et al. 2014), and when used in an antibody-drug conjugate (Pahl et al. 2018), alpha-amanitin successfully treated both drug-tolerant cancer cells and mice suffering cancer relapse (Kume et al. 2016).

For the last sixty years, commercial standards of the primary amatoxins (alpha, beta and gamma-amanitin) and primary phallotoxins (phalloidin, phallacidin) have been largely extracted from wild-foraged mushrooms, primarily from *Amanita phalloides* (Matinkhoo et al. 2018). Both phallotoxins and amatoxins are produced on the ribosome (Hallen et al. 2007), unlike many fungal chemical products, which are produced with nonribosomal peptide synthetases (NRPSs) (Bushley and Turgeon 2010). Though the genes encoding the cyclic peptide precursors are known (Hallen et al. 2007), the exact enzyme(s) responsible for the hydroxylation and epimerization processes remain elusive (Luo et al. 2018). In 2018, alpha-amanitin was synthesized chemically (Matinkhoo et al. 2018), but the process has yet to be scaled for mass production.

Before the effects of such compounds can be studied, the compounds must first be extracted and isolated. Procedures for extracting amatoxins from mushrooms have been evolving since the middle of the last century (Wieland et al. 1954). All of the amatoxin containing *Amanita* species are ectomycorrhizal, forming an obligate mutualism with trees, and cannot yet made to fruit in the lab (Smith and Read 2008). Mushroom fruiting is often seasonal, so the fruiting bodies must be collected over a relatively short period. A common protocol is to dry the mushrooms to prevent spoilage, allowing more time for extraction than work with fresh mushrooms, and then to grind the samples to a fine powder (Sgambelluri et al. 2014; Garcia et al. 2015). Most such studies have been performed on *Amanita phalloides*, due its global distribution and ample size. Previous extraction methods were often developed for preparing samples for instrumental or chromatographic detection. Most of these methods used solvent-based liquid extraction and required chromatography to ensure sufficient separation of other potentially interfering compounds within the sample, prior to detection by UV or mass spectrometry (MS).

Compared to MS, immunochemical (antibody-based) detection methods are less prone to interfering compounds but are not compatible with high amounts of solvents. Recently we developed immunochemical detection methods (ELISA and LFIA) for amatoxins (Bever et al. 2019; Bever et al. 2020). During this process, we adjusted the extraction procedure to be compatible for antibody-based detection, which meant removing the use of the organic solvent, methanol. For the purposes of making a rapid field portable detection method, we also determined sample maceration was not needed to extract the amatoxins, and that the incubation time for extraction could be accomplished in 1 minute.

In this current work, we hypothesized that these same extraction adjustments– minimal tissue maceration and reduced incubation time–could be transferable to instrumental (e.g., UV or mass spectrometry) detection methods. To test the need for sample maceration, we utilized three different techniques: 1) traditional mortar and pestle, 2) a similarly disruptive method of bead beating, and 3) no grinding, but rather hand shaking dried mushroom tissue in the extraction buffer. In addition, we performed the solvent extraction step at varying times to observe if more time allows for more toxin to be removed from the tissue. Lastly, we utilized two comparable solvent evaporation methods (rotovap or speedvac) to establish if multiple samples could be processed simultaneously, thus improving sample throughput.

## 1.3. Materials and methods

### 1.3.1. Extraction protocols

A single large, dried *Amanita phalloides* mushroom was selected for analysis. The mushroom was collected from Point Reyes National Seashore in 2017, under permit #PORE-2017-SCI-0054. Prior to extraction, the mushroom was re-dried at 113° F until it reached a constant weight.

The mushroom cap was radially divided into 16 pieces: it was first cut into four quadrants, then each quarter was further divided into four pieces. For the bead beating and hand shaken samples, pieces of dried mushroom were added to each vial, with a mass between 0.9 and 1.1 g. For the remaining samples, the rest of the mushroom cap was frozen with liquid nitrogen and ground with a mortar and pestle. Then, 0.9-1.1 g of mushroom powder was weighed into each 15 ml Falcon tube.

Bead beating was performed in 2.0 mL polypropylene screw cap vials (BioSpec Products, Inc., Bartlesville, OK, USA). Approximately 50 mg of 1.3 mm chrome steel beads (BioSpec Products) were used, and the sample was shaken on a Mini-Beadbeater 24 (BioSpec Products) for 2 minutes to pulverize the dry sample.

To each sample type, we then added 1 ml of extraction solution (80% methanol: 10% .01M HCL: 10% ddH_2_0) (Walton 2018) per 0.02 g of mushroom tissue. Samples were incubated for various incubation times; 1, 5, 10, 30 and 60 minutes. Samples that were only incubated for one minute were completed at room temperature. The samples that were incubated for 5, 10, 30 and 60 minutes were placed in a 30°C incubator and gently rocked for the allotted time.

Next, samples were centrifuged for 10 minutes at 4000 g, except for the hand shaken samples; for these, the volume was transferred to the next container. The samples were then rotary evaporated under vacuum with either a speedvac (temperature 45°C, Vacuum: 5) or rotovap (bath temperature 40°C). Rotovapped samples were evaporated in 100 mL round-bottom flasks.

Once the sample was completely dried, it was resuspended in 100 ul of LCMS-grade water per ml of extraction solvent used. The final samples were diluted (15 ul of sample, 50 ul of water) before being run on HPLC. Each treatment was repeated in duplicate.

### 1.3.2. HPLC

Extracts were analyzed on an Agilent 1200 series HPLC coupled to a UV detector. Compounds were separated over a Phenomenex Kinetex XB-C18 column (100 × 3 mm, 100 Å, 2.6 μm particle size) column held at 50× C using a gradient method with a mobile phase consisting of 20 mM ammonium acetate pH 5 (solvent A) and acetonitrile (solvent B). The gradient was as follows: 0-4 min. 6% B, 4-5 min. 6-15% B, 5-10 min. 15-18% B, 10-12 min. 18-60% B, 12-13 min. 60% B, 13-13.5 min. 60-6% B, 13.5-16 min. 6% B. Compounds were detected using a UV detector programmed to monitor wavelengths of 295 nm and 305 nm. Peaks were integrated in Agilent OpenLAB software. Alpha-, beta-and gamma-amanitin were quantified using the UV signal at 305 nm.

To generate estimates of each toxin concentration, dilutions (ranging from 12.5 - 250 ug/mL) of alpha-(Sigma, ≥90% purity), beta-(Sigma, ≥95% purity) and gamma-amanitin (≥90%, Enzo Life Sciences, Farmingdale, NY, USA) standards in LCMS-grade water were assessed. Linear regression analysis was performed using MassHunter software.

### 1.3.3. Data Analysis

All statistical analyses were carried out using Python SciPy. First, a test for normality was completed for each analyte. For those analytes with normally distributed data, t-tests were completed between each treatment type, while for non-normally distributed data sets, Mann-Whitney U tests were performed. The Bonferroni correction was applied to account for multiple comparisons. Comparisons were completed using the duplicate values from each evaporation type, between time points, and between the different methods used to macerate tissue samples.

## 1.4. Results

### 1.4.1. Overall toxin analysis

All samples in this study were analyzed using an HPLC-UV method. Liquid chromatography provided separation of the extracted components so that each compound could be detected independently. The retention times for alpha-, beta-, and gamma-amanitin were 3.821, 2.061, and 8.213 minutes, respectively. For statistical comparison between sample preparation methods, we utilized raw data (peak area). In every experimental condition tested, all three target compounds (a-AMA, b-AMA, g-AMA) were detected.

First a test of normality was performed on the data, which found that only the data for beta-amanitin was normally distributed. Kruskal Wallis tests were then run on the data from all three toxins, which found that extraction method was significant (p ⍰ 0.05 for each toxin), but found no significance due to drying method (p > 0.05 or time (p > 0.05) for any toxin.

Data from each toxin were then analyzed independently. For alpha-amanitin and gamma-amanitin, Mann-Whitney U tests were performed, but no differences were statistically significant.

### 1.4.2. Beta-amanitin

Because the beta-amanitin data were normally distributed, we investigated the data with t-tests followed by a Bonferroni correction for multiple comparisons.

For the samples that were dried with speedvac, hand shaking was not significantly different than any of the bead-beat samples (two-tailed t-tests, p > 0.05). There were also no statistical differences between hand shaking and mortar and pestle at any of the time points (two-tailed t-tests, p > .05).

For the samples that were dried with rotovap, hand shaking yielded significantly more toxin than bead beat samples incubated for every time point except for 1 minute (two-tailed t-tests, p < .05). There were no significant differences between hand shaking and mortar and pestle at any time points.

Comparing rotovap and speedvac, there were several significant differences between samples that were bead beat (two-tailed t-tests, p < .05). Beadbeat samples that were incubated for 10 minutes and rotovapped were significantly different from samples that were dried with speed vac and incubated for 30 minutes (p = .03) and 60 minutes (p = .03). Similarly, beadbeat samples that were incubated for 30 minutes and rotovapped were significantly different from samples that were dried with speed vac and incubated for 30 minutes (p < .05) and 60 minutes (p < .05). There was no statistical difference between samples that were hand-shaken (two-tailed t-test, p = 1.93). For samples that were ground with mortar and pestle, there was a significant difference between 30 minute rotovap and 30 minute speedvac samples (two-tailed t-test, p = .03).

## 1.5. Discussion

In this study, we extracted amatoxins using modified sample preparation methods. In many cases, previous studies focused on the development of analytical chemistry techniques, and not the wet lab steps leading to chemical analysis (Tanahashi et al. 2010; Nomura et al. 2012; Yoshioka et al. 2014; Zhang et al. 2016). Early protocols incubated for several days (Wieland and Wieland 1959), and then later for 24 hours or overnight (Mcknight et al. 2010; Clarke et al. 2012; Kaya et al. 2013). Only a handful of studies incubated for as short a period as an hour (Stijve and Seeger 1979; Jansson et al. 2012; Sgambelluri et al. 2014; Garcia et al. 2015). The shortest incubation step we could find in the literature was 10 minutes (Ahmed et al. 2010). In this study, we provide more evidence that reduced incubation times, down to as little as 1 minute, are sufficient to achieve toxin extraction from dried mushroom tissue.

At the outset of each extraction, most methods are performed on ground or macerated dried mushroom tissue (Sgambelluri et al. 2014; Garcia et al. 2015). This is both time-consuming and can generate airborne dust, which could increase exposure of the researcher and their environment to toxic materials. To reduce airborne dust production, we examined if bead beating, an equally destructive method for macerating tissue, self-contained within a tube, could be a suitable alternative. Our results indicate that the toxin is extracted through bead beating, although for rotovapped beta-amanitin samples, the amount of toxin extracted from bead beaten samples was significantly less than the amount of toxin extracted from handshaking (**Figures 1-3**). With bead beating, the higher variability and recovery loss was likely due to the volume of fluid trapped on and among the beads, thus reducing overall recovery. Recovery could be improved by washing the beads. However, the disruption to the cell wall does not seem needed, given how well the hand-shaken samples were extracted.

**Figure 1.**
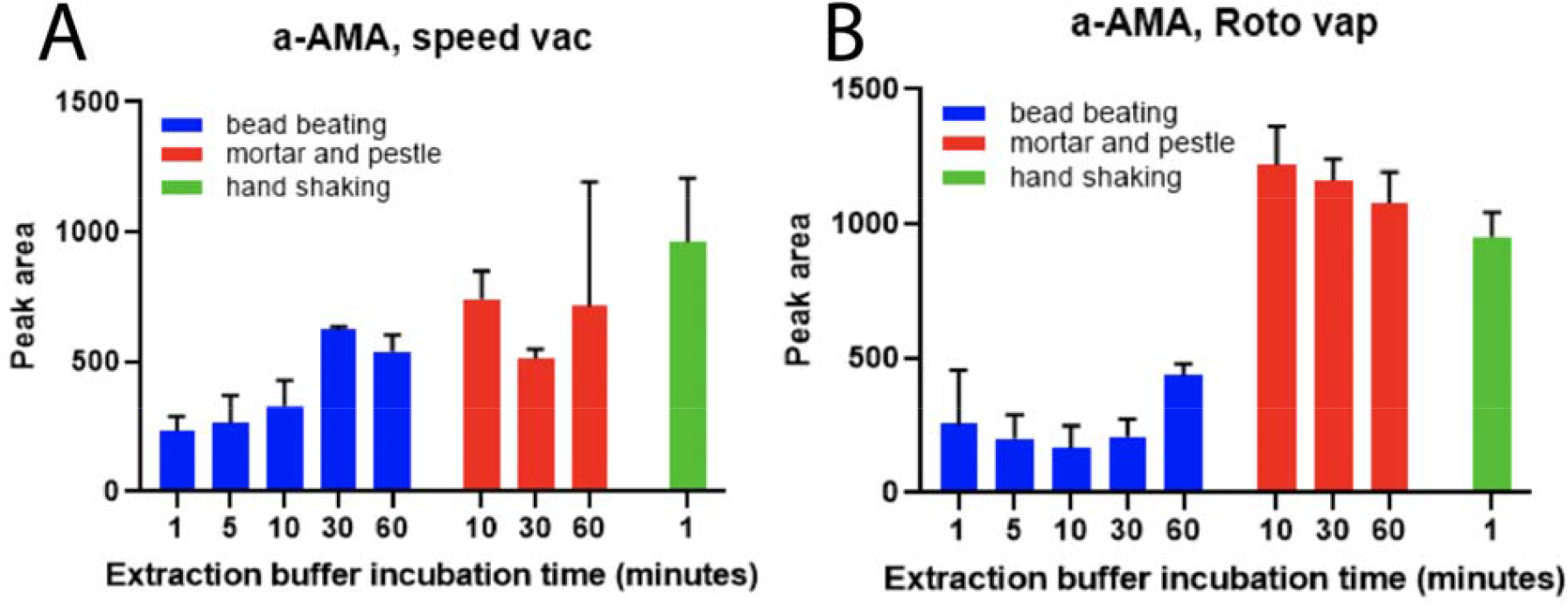
Extraction of Alpha-amanitin. Concentrations of Alpha-amanitin from mushroom extracted samples evaporated by (A) speed vac or (B) roto vap. Values are means +/-standard deviation (n=2). There were no statistically significant differences between any treatments.

As another alternative to reduce dust production, we examined the feasibility of extracting toxins by simply dropping a small piece of dried tissue in a tube and handshaking the tissue in extraction solvent. Our results indicate that hand shaking a piece of mushroom for a minute can yield as much toxin as a more elaborate protocol involving grinding the tissue with liquid nitrogen, followed by an hour-long incubation step (**Figures 1-3**). The only potential drawback of handshaking is that centrifuging is not as effective with mushroom pieces as it is with mushroom powder, and there is risk of small tissue transfer after the incubation step. However, this risk of tissue transfer could likely be reduced by performing a simple filtration of the extract.

This study is, to our knowledge, the first comparison of rotovap and speedvac on recovery of mushroom toxins, and perhaps the first such comparison of any biological compound. Some early studies on amatoxin extraction did not specify what type of equipment was used to evaporate solvent, instead simply stating that the solvent was evaporated under vacuum (Faulstich et al. 1973; Stijve and Seeger 1979). Our results indicate that, for alpha-amanitin and gamma-amanitin, evaporating multiple samples simultaneously with a speed-vac yielded as much toxin as evaporating samples one at a time with a rotovap, but the same was not always true for beta-amanitin (**Figure 2**). If the primary goal is to maximize the amount of amatoxin extracted, then rotovap may be more appropriate. We think that, for beta-amanitin, the yield with rotovap is higher than speedvac because, with the speedvac and running multiple samples simultaneously, the extract can sometimes over-dry to the walls of the container and fail to re-dissolve. This over-drying may occur with these mushroom toxins as well as many other biological compounds, warranting further investigation. Future work could also consider comparing rotovap and speedvac to evaporating solvent under a liquid nitrogen stream (Clarke et al. 2012).

**Figure 2.**
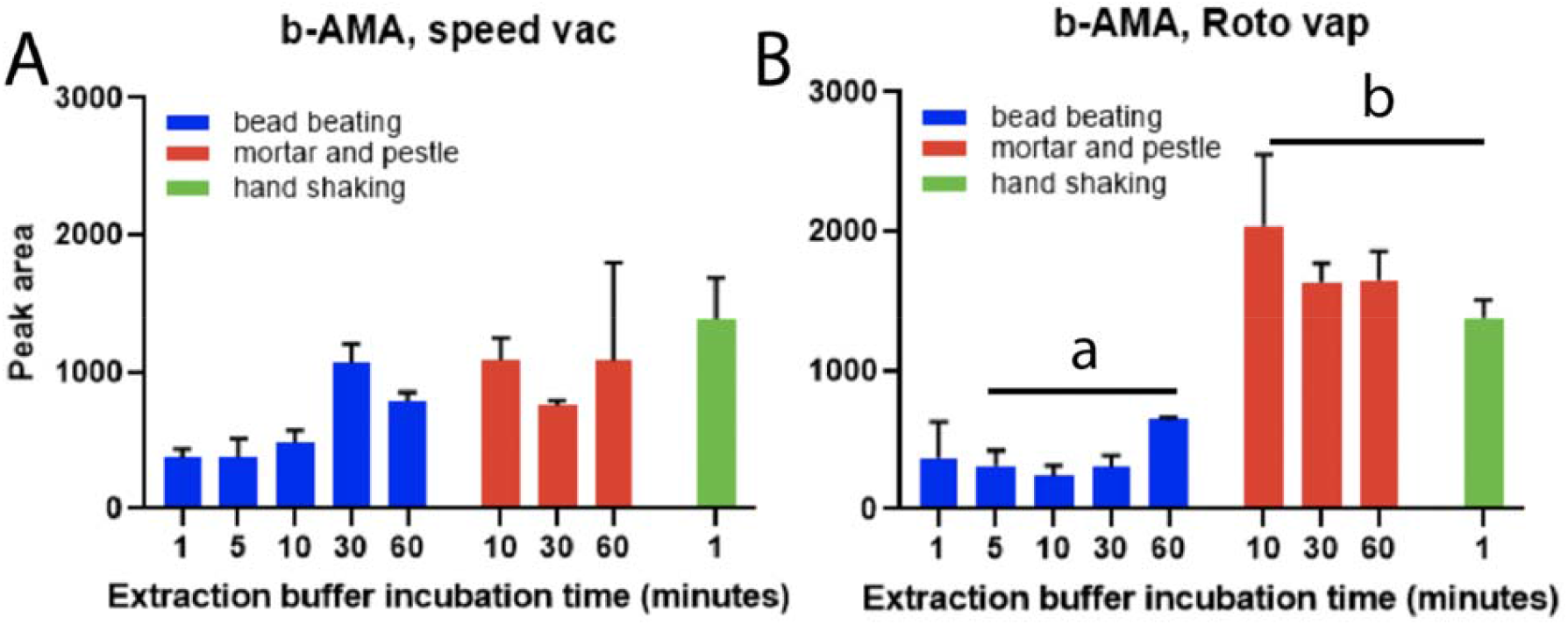
Extraction of Beta-amanitin. Concentrations of Beta-amanitin from mushroom extracted samples evaporated by (A) speedvac or (B) rotovap. Values are means +/-standard deviation (n=2).

**Figure 2.**
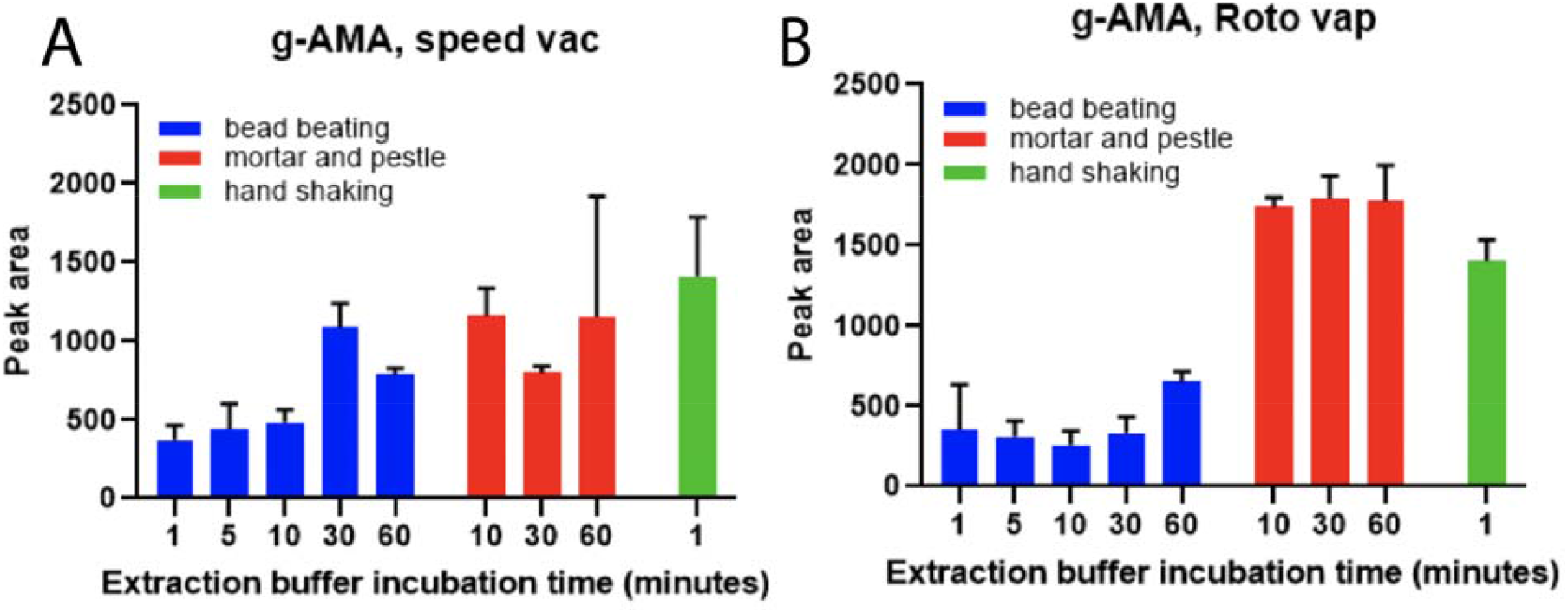
Extraction of Gamma-amanitin. Concentrations of Gamma-amanitin in mushroom extracted samples evaporated by (A) speedvac or (B) rotovap. Values are means +/-standard deviation (n=2).

Together, our results allow us to theorize about the cellular nature of the toxin within the mushroom. Because hand shaking was always as efficient as mortar and pestle at extracting amatoxins, we can surmise that amatoxins are not deeply embedded in cell walls or compartments, and thus do not need extensive pulverization for extraction. This observation that the amatoxins are not deeply embedded allows us to postulate about the ecological role of amatoxins. Such readily released toxins may be more accessible to fungivores such as *Amanita*-associated *Drosophila* species (Greenleaf et al. 1979; Stump et al. 2011; Mitchell et al. 2017), and thus act as a potent deterrent against mycophagy.

The ease of extraction can also help to explain the bioavailability of the amatoxin. For animals, the toxicity of amatoxins is directly related to the ability of the species in question to absorb the toxins in the gut (Wieland and Faulstich 1978). As recently as this last decade, it has been erroneously stated that the toxins were not water soluble (Allen et al. 2012), and this idea has been perpetuated throughout some mycological communities. Our work suggests that even the commonly practiced method of ‘taste testing’ a small piece of mushroom would yield toxin exposure to a person and thus is not recommended when one suspects the mushroom to be a species that contains amatoxins. Furthermore, perhaps the toxin is produced in such relative excess because it can readily leave the mushroom with water, with precipitation such as rain or fog drip. Once in the soil, amatoxins could function not only as a defense chemical but perhaps also an offense chemical, inhibiting nearby fungi and other eukaryotic soil microbes.

In conclusion, we present an extraction protocol that saves time, reduces equipment contamination, and minimizes risk to the researcher. Using liquid nitrogen and grinding the mushroom to a fine powder poses potential harm to the researcher and potentially contaminates tools and the surrounding environment. In contrast, handshaking a piece of mushroom reduces occupational hazards for the scientists, allowing them to reduce both specimen handling time and destructive manipulation methods. Toxicologists and officials should be aware of potential nefarious acts to endanger pets (dogs and cats) and humans, considering the ease of toxin extraction with minimal equipment and technical experience, and that this mushroom species is still undergoing a range expansion (Wolfe and Pringle 2012). Furthermore, the impact of this faster, safer method may help produce these important toxins faster, for both research and medical use.

## 1.6. Acknowledgements

The authors wish to thank Jacob Baker, Gaby Olea Vargas, and Stephen Shen for their assistance with protocol development. Sincere gratitude to the Traxler lab for use of their speedvac and rotovap, and to Michael Filigenzi at UC Davis for sample processing. Thank you to the Bruns and Taylor labs for feedback on an early version of the manuscript. Finally, thank you to the late Jonathan Walton for the gift of his original extraction protocol, as well as his enthusiasm and insight.

